# Deep learning models for chemical perturbation prediction do not yet utilise drug molecular features

**DOI:** 10.64898/2026.05.13.724458

**Authors:** Jinming Bai, Sharon Prince, Geoff S. Nitschke

**Author notes:** Contributing author.

## Abstract

Recent deep learning models for L1000 chemical perturbation prediction incorporate dedicated drug molecular encoders. We retrained seven such models from scratch with zeroed or shuffled drug inputs, and compared them with a multilayer perceptron that uses only cell-line basal expression. Under drug-blind evaluation, ablation caused negligible performance changes and the drug-free baseline matched all models. Current architectures do not yet utilise drug molecular features for generalisation to unseen compounds.

---

The large-scale L1000 chemical perturbation transcriptomic data generated by the LINCS programme has given rise to a series of deep learning models that aim to predict drug-specific gene expression changes from molecular structure. Since 2021, seven such models have been published—DeepCE[1], CIGER[2], MultiDCP[3], TranSiGen[4], PRnet[5], PertDiT[6] and XPert[7]—each with a dedicated drug encoding module, spanning graph convolutional networks[8], variational autoencoders[9], diffusion Transformers[10] and dual-branch attention architectures[11] (Extended Data Fig. 1). The core premise of these models is that the input drug molecular features will be utilised by the model to generate drug-specific transcriptomic predictions.

In the present study, we performed drug molecular feature ablation on all seven models (Extended Data Fig. 1) to assess whether they actually utilise their drug inputs. We also introduced two naive baselines—a global mean (Mean) and a per-cell-line mean (Mean cell)—as well as a deliberately simple three-layer MLP that takes only cell-line basal expression as input and receives no drug information, to quantify the predictive gain actually conferred by drug molecular features in current models. Three of the models (DeepCE, CIGER, MultiDCP) were not originally designed to accept sample-paired L1000 basal expression profiles; in the figures, we marked their results with a dagger (†).

To ensure a fair comparison, we deployed all seven models on the same L1000 single-dose single-time-point dataset released by XPert (978 landmark genes, approximately 78,000 samples), using drug-blind five-fold cross-validation such that test drugs were entirely absent from the training set. As the models required adaptation to a unified data format, we first verified that the engineering modifications did not affect their original performance: all models under the full drug input (Full) condition matched the values reported in their respective publications or those obtained from running the original codebases (Extended Data Fig. 2).

We designed two ablation conditions: (1) Zero, in which drug features were replaced with an all-zero vector; and (2) Shuffle, in which drug features were randomly permuted across samples once before training and then held fixed, preserving the feature distribution but breaking drug–sample pairing. All models were retrained from scratch under both ablation conditions. The Mean baseline always predicts the global average differential expression across the training set; Mean cell predicts the per-cell-line average; neither uses drug information. The MLP was selected via hyperparameter sweep (Extended Data Fig. 3) as a three-layer fully connected network (hidden dimension 2,048), taking only the 978-dimensional basal expression as input.

The difference in differential-expression Pearson correlation (PCC_DEG_) between the Full and Zero conditions was negligible for all seven models: the largest absolute difference was only 0.012 (Fig. 1a). The shift under Shuffle was slightly larger (maximum 0.027). We also examined 31 evaluation metrics, including Spearman correlation, *R*^2^, Precision@*K*, Wasserstein distance and DEG-weighted calibration metrics, all of which converged on the same conclusion (Extended Data Fig. 4, 5, 6). Repeating the ablation experiment on each model’s original dataset and original splitting scheme produced consistent results (Extended Data Fig. 7).

**Figure 1:**
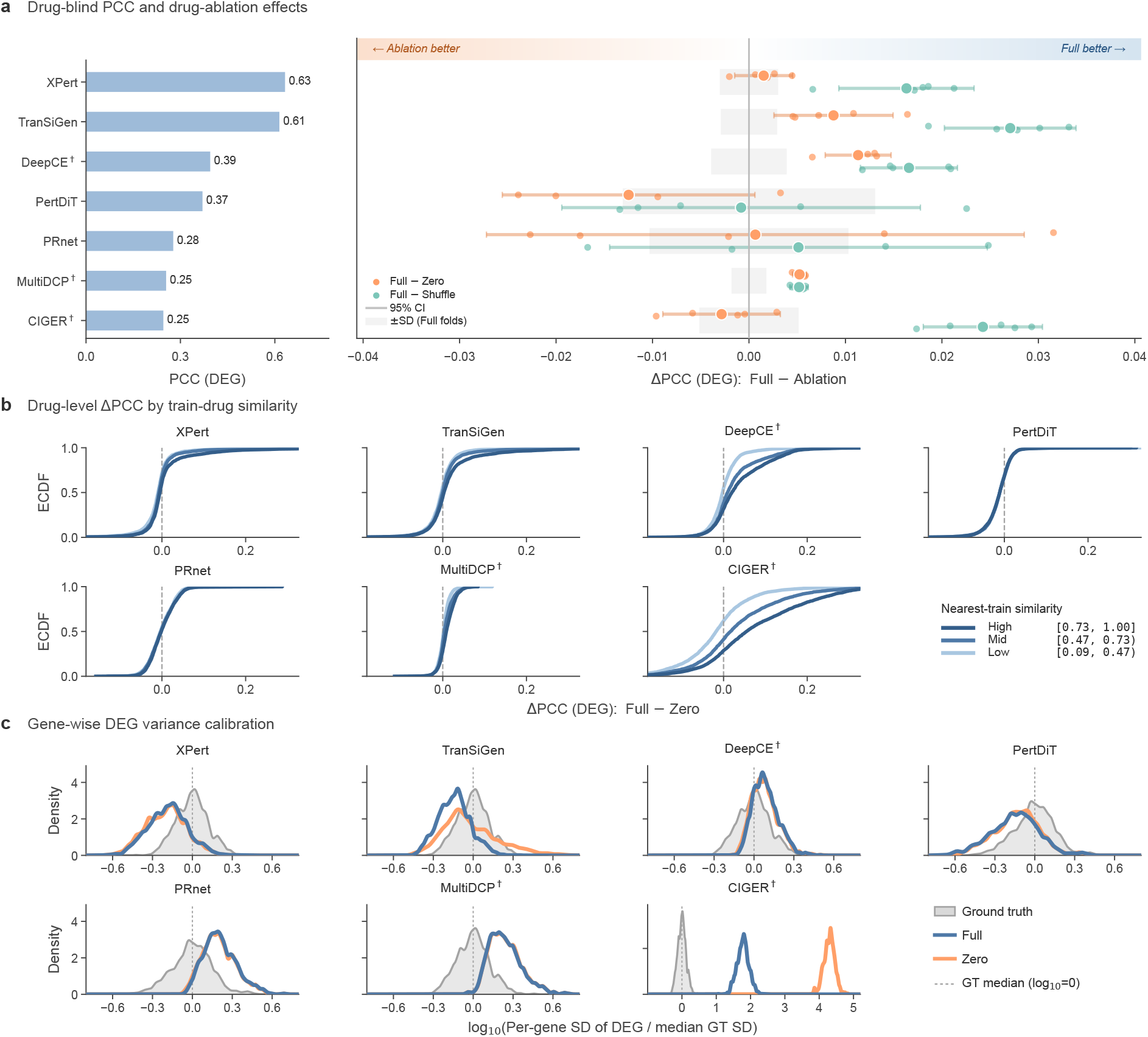
Drug-input ablation under drug-blind evaluation. **a**, Drug-blind PCC (DEG) and paired drug-ablation effects for the seven published models. Left, Full-condition PCC (DEG). Right, paired effect sizes computed as ΔPCC (DEG) = Full − Zero and Full − Shuffle. Points indicate fold-level estimates, larger points indicate across-fold means and horizontal segments indicate 95% CIs across five folds (Student’s *t*, df = 4). Grey bands indicate *±* s.d. across Full-condition folds. **b**, Empirical cumulative distribution functions of per-drug ΔPCC (DEG) = Full − Zero, stratified by nearest-train compound similarity. **c**, Gene-wise DEG variance calibration, shown as the distribution of log_10_(per-gene s.d. of predicted DEG /median ground-truth s.d.). The grey distribution denotes the ground truth, and coloured curves denote Full and Zero conditions. Dashed vertical lines mark the ground-truth median. † Models not originally designed to accept sample-paired L1000 basal expression profiles.

We further ruled out the possibility that the ablation effect was masked by averaging across chemical space. When test drugs were stratified by Tanimoto similarity to the nearest training compound into high, mid and low groups, the cumulative distributions of ΔPCC overlapped closely for most models (Fig. 1b). Using a scaffold split to partition training and test sets by molecular scaffold, the same conclusion held (Extended Data Fig. 7). At the cell-line level, ΔPCC was overall close to zero across most models, with no stable global positive gain; variation in ΔPCC was strongly model- and cell-line-dependent, and positive shifts were confined to subsets of cell lines and models (Extended Data Fig. 8).

We also examined the effect of ablation on the variance structure of predictions. For the six non-CIGER models, the per-gene prediction standard deviation distributions under Full and Zero were nearly identical (Fig. 1c, Extended Data Fig. 9), indicating that the drug input did not introduce drug-specific predictive variation.

Next, we assessed whether a simple model using only cell-line basal expression could achieve the same performance. The drug-free MLP achieved a drug-blind PCC_DEG_ of 0.637, comparable to XPert (0.633), the strongest of the seven full-feature models, and far exceeding both Mean (0.099) and Mean cell (0.243) (Fig. 2a). This reveals a clear information hierarchy: Mean *<* Mean cell *<* MLP *≈* Full models—the primary source of predictive power is cell-line basal expression, and drug features do not confer additional benefit.

**Figure 2:**
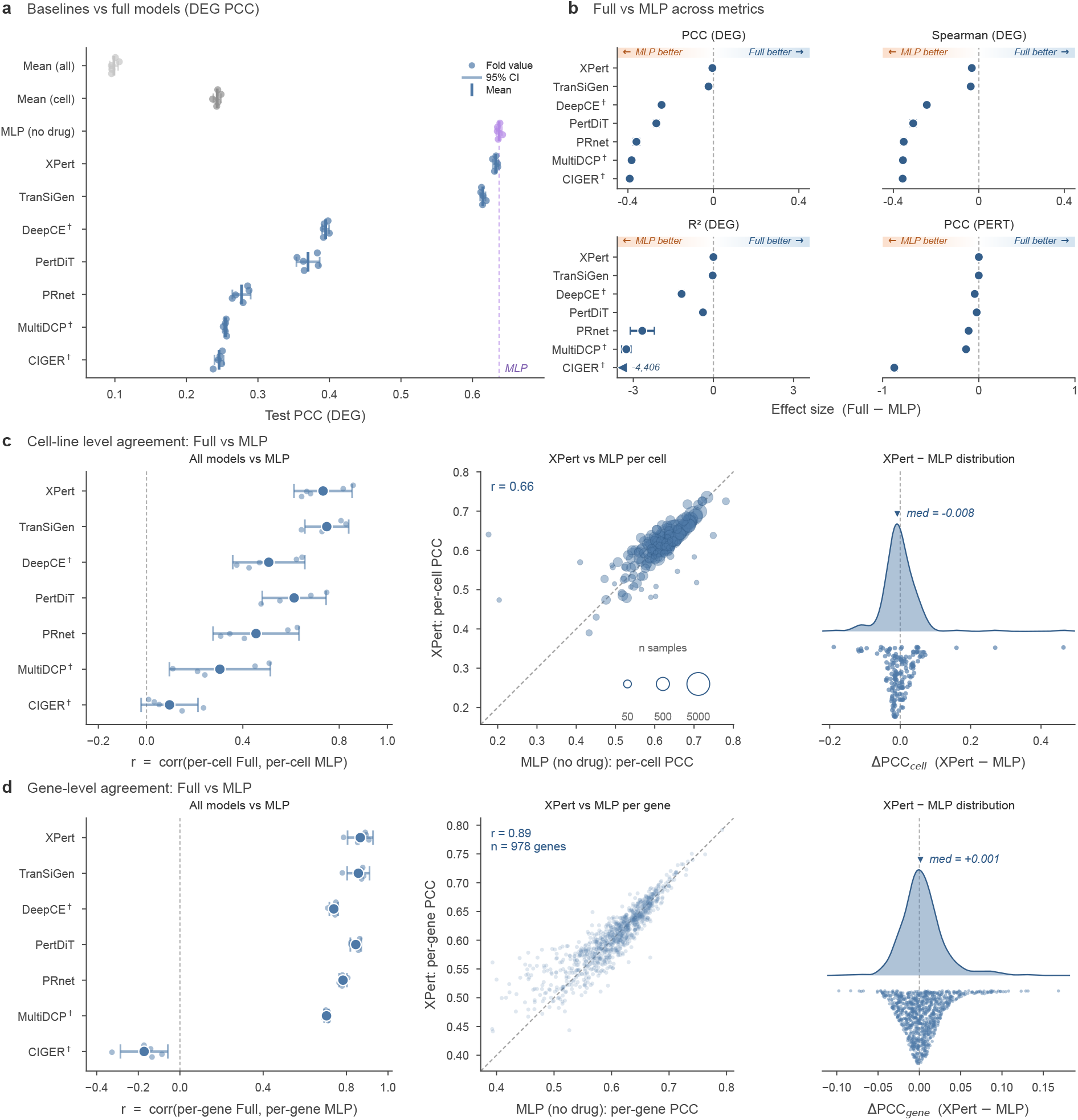
Comparison between Full models and the MLP (no drug) baseline. **a**, Test PCC (DEG) of mean baselines, the MLP (no drug) and the seven published models under drug-blind evaluation. Points indicate fold-level estimates and horizontal segments indicate 95% CIs across five folds (Student’s *t*, df = 4). **b**, Effect sizes comparing each Full model with the MLP baseline across PCC (DEG), Spearman (DEG), *R*^2^ (DEG) and PCC (PERT). Positive values favour the Full model, as indicated by the direction bands. **c**, Cell-line level agreement between Full models and the MLP. Left, correlation between per-cell Full and per-cell MLP PCC. Middle, XPert versus MLP per-cell PCC; bubble area is proportional to the number of samples. Right, distribution of per-cell ΔPCC, computed as XPert − MLP. **d**, Gene-level agreement between Full models and the MLP. Left, correlation between per-gene Full and per-gene MLP PCC. Middle, XPert versus MLP per-gene PCC. Right, distribution of per-gene ΔPCC, computed as XPert − MLP. † Models not originally designed to accept sample-paired L1000 basal expression profiles.

Across representative metrics, Δ(Full − MLP) was mostly centred around zero, and no model consistently outperformed the MLP across all metrics (Fig. 2b, Extended Data Fig. 4). At the cell-line level, per-cell PCC values of each model correlated with those of the MLP (*r*_cell_ range 0.31–0.75), with higher-performing models showing stronger agreement. For XPert, the per-cell PCC scatter against the MLP was linear (*r* = 0.66), with a median difference of only −0.008 (Fig. 2c). At the gene level, the agreement was even more pronounced: *r*_gene_ ranged from 0.70 to 0.89 (excluding CIGER), and XPert versus MLP per-gene PCC values fell nearly on the diagonal (*r* = 0.89; Fig. 2d).

In summary, seven models spanning graph convolutional networks, variational autoencoders, diffusion Transformers and dual-branch attention architectures all showed negligible contribution of drug molecular features under drug-blind evaluation, despite their substantially higher computational cost compared to the drug-free MLP (Extended Data Fig. 10).

The asymmetry between Full *≈* Zero and Shuffle performing slightly worse is itself diagnostically informative. It indicates that the drug branch is indeed partially read by the models—incorrect drug inputs cause greater harm than no drug input—but this does not mean the models effectively utilise correct drug features. With the drug branch removed, models can still rely on the cell-line expression backbone to maintain performance; when presented with incorrect but statistically plausible drug inputs, sample-specific erroneous modulation occurs. Crucially, the magnitude of this damage (*<* 0.03 PCC) is far smaller than the performance gap one would expect if the drug channel truly carried effective chemical information, indicating that the utilisation of drug features is fragile and limited—a sensitivity to incorrect conditioning rather than a robust dependence on correct conditioning.

This conclusion is consistent with recent findings in adjacent fields. In genetic perturbation prediction, Ahlmann-Eltze *et al*.[12] found that single-cell foundation models did not outperform simple linear baselines; in drug sensitivity prediction, Branson *et al*.[13] reported that deep learning models failed to surpass baselines that did not use molecular features. Our work extends this finding to chemical perturbation transcriptomic prediction.

Our benchmark has several limitations. All experiments were conducted under L1000 single-dose single-time-point conditions (10 *µ*M, 24 h), and multi-dose scenarios were not covered. All cell lines are cancer cell lines. Some models were not originally designed to accept sample-paired L1000 basal expression profiles, but the ablation comparison is an internal control within each model and is unaffected by this.

Our results do not negate the potential value of drug molecular information in principle, but indicate that the model architectures and training procedures evaluated here have not yet effectively utilised this information to achieve generalised prediction for unseen drugs. We expect that rigorous input-utilisation validation—such as the ablation experiments and drug-free baselines employed here—will help drive substantive progress in this field.

## Methods

### Data

We used the L1000 single-dose single-time-point subset released with XPert[7], comprising 78,453 paired perturbation observations from 8,276 compounds across 164 cell lines. All perturbations were measured at 10 *µ*M for 24 h. Each observation contained a perturbed expression profile, *x*_pert_, and its matched basal expression profile, *x*_ctrl_. All analyses were restricted to the 978 L1000 landmark genes.

For each sample, the prediction target was the differential-expression vector

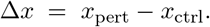

Models that produced absolute perturbed expression profiles were converted to differential-expression predictions by subtracting the corresponding *x*_ctrl_. Models that natively predicted differential expression were evaluated directly. Unless otherwise stated, all metrics were computed on Δ*x*.

### Models and adaptation

We evaluated seven published L1000 chemical perturbation prediction models at pinned upstream commits: DeepCE[1] (`9b0d04f`), CIGER[2] (`81c16f1`), MultiDCP[3] (`36ecdef`), TranSiGen[4] (`8ec2218`), PRnet[5] (`f19174b`), PertDiT[6] (`596d681`) and XPert[7] (`d53ff49`). For each model, we wrote a training shim that imported the upstream model code and harmonised model inputs and outputs to a common interface, including drug-blind split labels and differential-expression predictions. Unless stated below, hyperparameters, optimisers, schedulers and architectural defaults were kept as in each model’s original publication or released configuration.

DeepCE and CIGER do not accept sample-paired L1000 basal expression profiles; cell-line context was therefore supplied as a one-hot index. CIGER’s original 1,107-gene read-out was remapped to the 978 L1000 landmark genes through HGNC alias updates. MultiDCP supplied cell-line context through its `cell transformer` branch, which expects a basal-expression vector; same-cell within-drug technical replicates were collapsed to the median-norm sample, following the upstream pipeline. For TranSiGen, the two upstream shRNA-VAE checkpoints were loaded as initialisation for the joint VAE1/VAE2 model, which was then trained end-to-end with the upstream KPGT 2,304-d drug embedding. For PRnet, an *O*(*N*) `list.index()` call in the upstream data loader was replaced with a dictionary lookup as a semantics-preserving performance fix; the model retained its 1,024-bit FCFP4 fingerprint input. For PertDiT, MolT5, BioLinkBERT and per-dose embeddings were precomputed once and reused across folds. XPert was trained from the upstream `config l1000.yaml` with both drug inputs intact: the UniMol per-atom tensor and the heterogeneous-graph drug embedding. Per-file SHA-256 checksums against the pinned upstream commits and line-level diffs for modified files are provided with the released code.

To verify that harmonisation did not degrade model performance, we additionally retrained each model on its original-paper dataset, split and primary metric, and compared the reproduced values with the published or original-code values. Because the original studies used different datasets, splits and metrics, this reproduction analysis was used only to assess implementation fidelity, not for cross-model ranking.

### Drug-blind cross-validation

We generated five drug-blind folds by randomly partitioning the 8,276 unique compounds into five groups using seed 131419. For fold *i*, group *i* served as the test set, group (*i* + 1) mod 5 as the validation set and the remaining three groups as the training set, yielding a 3:1:1 train–validation–test ratio at the compound level. All samples from the same compound were assigned to the same split, ensuring that test compounds were absent from both the training and validation sets.

All seven models were trained from scratch on each fold under the Full condition and under the ablation conditions described below. Model selection used each model’s existing validation convention, and test predictions were generated only from the selected checkpoint.

### Drug-feature ablation

We defined two ablation conditions to test whether model predictions depended on correctly paired molecular features. In both conditions, sample labels, cell-line inputs, basal expression profiles and expression targets were unchanged. All models were retrained from scratch under each ablation condition using the same split and hyperparameter settings as in the Full condition.

In the *Zero* condition, the drug-specific representation was replaced with zeros at the earliest point at which it could be cleanly excised before being combined with cell-line or basal-expression inputs. The intervention site was the NeuralFingerprint output for DeepCE, CIGER and MultiDCP; the FCFP4 input tensor for PRnet; the KPGT feature embedding for TranSiGen; the per-drug embedding entries for PertDiT, excluding the negative-control token while preserving dose information; and both the UniMol atom tensor and heterogeneous-graph drug embedding for XPert. For PRnet, we additionally clamped the CombAdaptor linear bias at zero with `requires grad=False`, because otherwise the bias term could absorb a learnable drug-independent constant when the drug input was zero.

In the *Shuffle* condition, drug identities were globally permuted across compounds using seed 131419 before training and held fixed for all epochs. For a sample with compound *d*, the model received the molecular feature of *π*(*d*), where *π* denotes the fixed permutation over unique compounds. This preserved the marginal distribution of molecular features while breaking the drug–sample correspondence. For XPert, the same permutation was applied to the precomputed UniMol and heterogeneous-graph arrays; for the other models, the permutation was applied at the data layer immediately before model input.

The Full–Zero comparison tested whether removing compound-specific information changed model performance. The Full–Shuffle comparison tested whether statistically plausible but incorrectly paired molecular features perturbed model predictions.

## Drug-free baselines

We included three baselines that received no drug-specific input.

The *Mean* baseline predicted the global mean of training-set differential-expression vectors for every test sample,

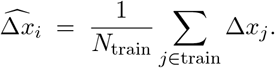

The *Mean cell* baseline predicted the mean differential-expression vector of the corresponding cell line in the training set,

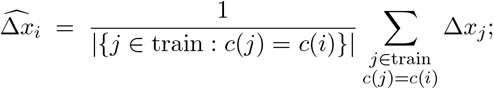

if a test cell line was absent from the training set, the global training mean was used.

The *drug-blind MLP* baseline was a feedforward network with three hidden layers of 2,048 units each, ReLU activation and dropout *p* = 0.1. The network received only *x*_ctrl_ as input and predicted Δ*x*; the final perturbed-expression prediction was

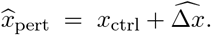

The MLP was trained with Adam, learning rate 10^−4^, batch size 512, gradient clipping with maximum norm 1.0 and early stopping with patience 50 evaluations on validation PCC_DEG_. The loss was

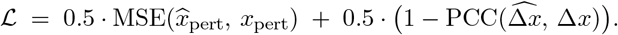

The default architecture was selected by an exhaustive grid sweep over depth *L* ∈ {1, 2, 3, 4} and width *H* ∈ {256, 512, 1024, 2048, 4096}, evaluated across all five drug-blind folds. We selected *L* = 3 and *H* = 2,048 as the smallest configuration at the saturation point.

### Evaluation

The primary metric was per-sample Pearson correlation on differential expression,

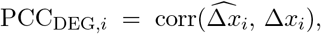

computed across the 978 landmark genes. Fold-level PCC_DEG_ was the mean across test samples.

We evaluated a unified panel of 31 metric entries covering correlation, goodness of fit, error, distributional distance, directional retrieval and high-signal DEG-focused performance. Correlation metrics included Pearson and Spearman correlations on Δ*x* and *x*_pert_, as well as per-drug PCC_DEG_. Goodness-of-fit metrics included per-gene *R*^2^, computed across samples for each gene and averaged across genes. Error metrics included MSE, RMSE, MAE and *L*_2_ distance. Distributional metrics included per-gene Wasserstein distance and RBF-kernel maximum mean discrepancy with *γ* = 1.0. Directional retrieval metrics included Precision@*K* for up- and down-regulated genes with *K* ∈ {10, 20, 50, 100}.

Following Miller *et al*.[14], we included high-signal and weighted metrics to assess performance on genes with stronger transcriptional responses. These included PCC and *R*^2^ restricted to the top 50, 100 and 200 genes ranked by mean |Δ*x*|, weighted MSE and weighted *R*^2^ using per-sample gene weights |Δ*x*|, and Precision@*K* metrics in which the reference set was defined as the top or bottom 100 genes of the observed Δ*x*, and the predicted set as the top or bottom *K* genes of the predicted Δ*x*.

For per-drug analysis, PCC_DEG_ was first computed for each test sample and then averaged over samples from the same compound. Cell-line-level analysis used the analogous aggregation by cell line. Gene-level analysis computed Pearson correlation across test samples separately for each gene.

Paired effect sizes were computed within fold as

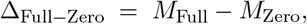

and analogously for Full–Shuffle and Full–MLP comparisons, where *M* denotes a given metric. Positive values indicate better performance under the Full condition for correlation and fit metrics; for error and distance metrics, the favourable direction is reversed. Across-fold means are reported with 95% CIs (Student’s *t* distribution, df = 4, *n* = 5 folds).

### Chemical-similarity and scaffold analyses

Per-drug performance was defined as the mean PCC_DEG_ across test samples from the same compound. Chemical similarity was quantified as the maximum Tanimoto coefficient between each test compound and any training compound in the same fold, using 2,048-bit Morgan fingerprints with radius 2 computed with RDKit[15]. Test compounds were stratified into three equal-frequency similarity bins, denoted high, mid and low, for empirical cumulative distribution analyses.

For the scaffold analysis, compounds were grouped by exact Bemis–Murcko scaffold using RDKit[15]. Acyclic compounds were pooled into a single acyclic bucket, and SMILES strings that failed to parse were pooled into a separate invalid bucket. Scaffold groups were assigned to five folds using a longest-processing-time greedy heuristic that sorted groups by sample count and assigned each group to the currently least-loaded fold, with seed 131419 and a maximum cross-fold sample-count deviation below 15%. The same 3:1:1 fold rotation used for the random drug-blind split was then applied. The split generator verified that no scaffold was shared between training, validation and test sets within any fold.

### Variance-calibration analysis

To assess whether drug inputs introduced drug-specific predictive variation, we computed the per-gene standard deviation of predicted differential expression across test samples and compared the resulting distributions with the ground truth under Full and ablated conditions. For visualisation, each per-gene standard deviation was normalised by the median ground-truth standard deviation and shown on a log_10_ scale. The Wasserstein distance between predicted and ground-truth gene-wise standard-deviation distributions was used as a single-number summary of variance mismatch.

### Original-setting ablations

As a robustness check, we repeated the Zero ablation under each model’s original data setting where applicable. Six of the seven models were retrained on their own published dataset and split convention, using the same intervention rule as in the unified benchmark: the drug-specific representation was zeroed immediately before it entered the cell-expression pathway. XPert was not run as a separate original-setting task because its original single-dose single-time-point dataset is the same dataset used for the unified benchmark.

### Computational cost

For each training run, the pipeline recorded wall-clock time, number of epochs run, selected epoch, number of trainable PyTorch parameters and test-set size. To compare models under a common hardware protocol, we re-measured per-iteration training time and peak GPU memory on a single NVIDIA A100-40GB GPU using each model’s original code, with 60 training iterations per run, including 10 warmup iterations and 50 timed iterations. Training speed is reported as the mean time per optimisation iteration, and model size is reported as the number of trainable parameters. These measurements were used only to contextualise predictive performance and were not used for model selection.

### Software

Experiments were implemented in Python 3.9 with PyTorch 2.1.0 and CUDA 12.1 in a single conda environment. RDKit 2024.3.2 was used for molecular fingerprints and Bemis–Murcko scaffold extraction. Data processing and evaluation used NumPy 1.26.4, SciPy 1.13.1, pandas 2.3.0, scikit-learn 1.6.1 and Scanpy 1.9.8. The full pinned environment is provided with the released code. Claude Code (Anthropic) and ChatGPT (OpenAI) were used as programming assistants for selected coding and debugging tasks. All AI-assisted code was reviewed, modified and tested by the authors before use.

## Supplementary information

Source Data files for Fig. 1, Fig. 2 and Extended Data Figs. 2–10 are provided with this manuscript.

## Acknowledgements

Computations were performed using facilities provided by the University of Cape Town’s ICTS High Performance Computing team (hpc.uct.ac.za; https://doi.org/10.5281/zenodo.10021612) and by the Centre for High Performance Computing (CHPC), South Africa, under programme HEAL1643. J.B. thanks Jan Buys (Department of Computer Science, University of Cape Town) for contributing GPU resources to the UCT HPC facility used in this study.

## Declarations

### Data availability

The L1000 single-dose single-time-point dataset used for the unified benchmark was obtained from the XPert[7] data release (figshare: https://doi.org/10.6084/m9.figshare.28955141; Zenodo: https://doi.org/10.5281/zenodo.15357711). Upstream datasets used for original-setting reproduction were obtained from the corresponding model deposits or public archives. Trained model weights, per-fold predictions and per-fold evaluation metrics for all models and ablation conditions are deposited at https://zenodo.org/records/20081274. Source data for all main and Extended Data figures are provided with the paper.

### Code availability

Code to reproduce data preparation, model training, drug-feature ablation, unified evaluation and figure-level source data generation is available at https://github.com/baijinming97/drug-perturbation-benchmark.

### Author contributions

J.B., G.S.N. and S.P. conceptualised the study. J.B. developed the methodology, implemented the software, performed the formal analysis and investigation, curated the data, prepared the visualisations and wrote the original draft. G.S.N. and S.P. supervised the project, acquired funding and reviewed and edited the manuscript. All authors read and approved the final manuscript.

## Extended Data

**Extended Data Fig. 1:**
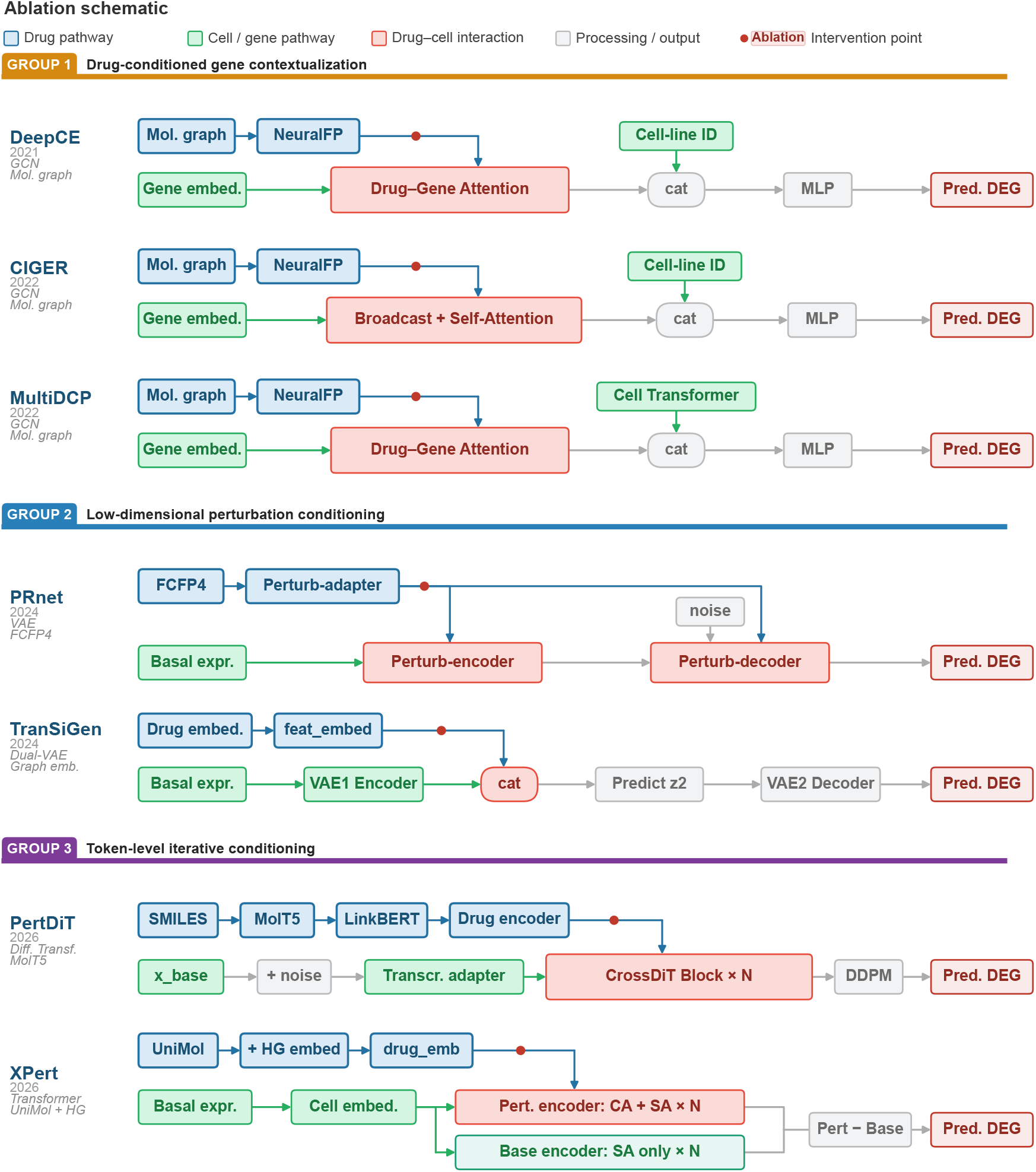
Ablation schematic across evaluated models. Schematic of the seven evaluated models, grouped by the stage at which compound information is integrated with cell-line and gene representations. Group 1 includes drug-conditioned gene contextualisation models; Group 2 includes low-dimensional perturbation-conditioning models; and Group 3 includes token-level iterative-conditioning models. Blue denotes the drug pathway, green the cell or gene pathway, red drug–cell interaction modules and grey processing or output components. Red dots indicate the architectural intervention points at which drug-specific representations were set to zero in the Zero ablation.

**Extended Data Fig. 2:**
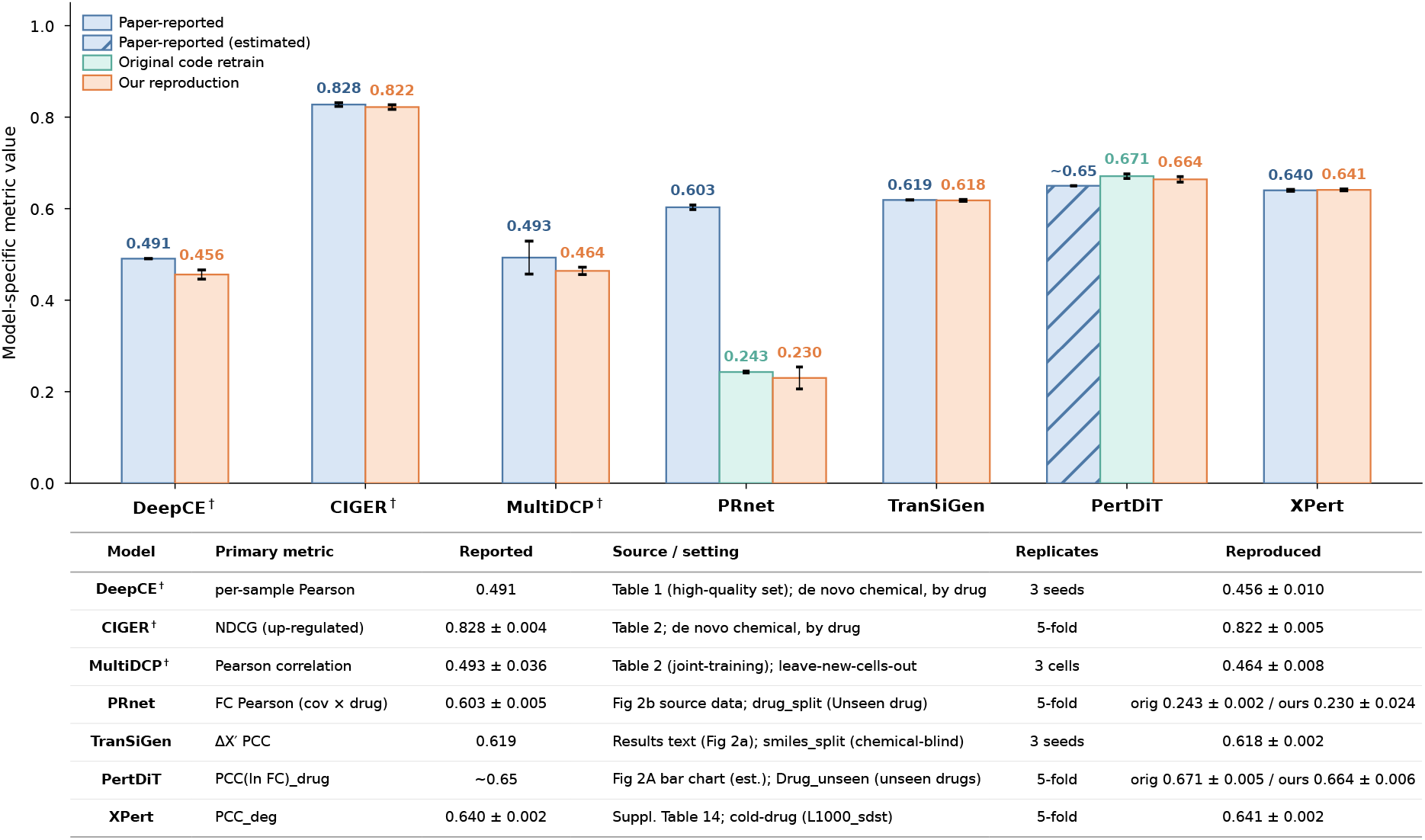
Reproduction fidelity of paper-reported metrics. Comparison between the primary performance metric reported in each model’s original publication and our reproduction of that metric under the corresponding source task or harmonised drug-blind evaluation where applicable. For models with available released code and original evaluation splits, original-code retrain results are shown separately. Bars show across-replicate means and error bars indicate s.d. across folds, seeds or cell splits, as specified in the table. Hatched bars denote paper-reported values reverse-estimated from published figures when no exact tabulated value was available. Each model is compared using its own reported primary metric and values are not intended for direct cross-model comparison. † Models not originally designed to accept sample-paired L1000 basal expression profiles.

**Extended Data Fig. 3:**
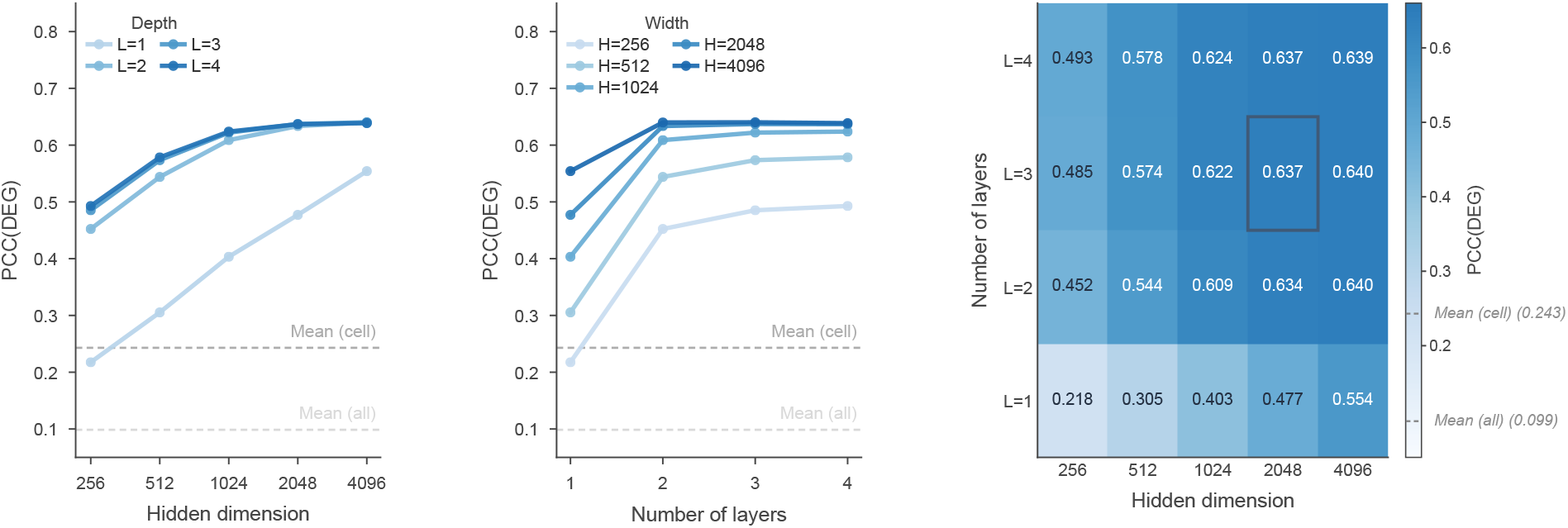
MLP hyperparameter sweep across depth and width. **a**, PCC (DEG) of the MLP (no drug) as a function of hidden dimension, stratified by network depth. Each point denotes the across-fold mean. Dashed horizontal lines indicate the Mean (all) and Mean (cell) baselines. **b**, PCC (DEG) as a function of network depth, stratified by hidden dimension. **c**, Heatmap of PCC (DEG) across hidden dimension and number of layers. The boxed cell indicates the default MLP configuration used in the main analysis, with three hidden layers and 2,048 hidden units.

**Extended Data Fig. 4:**
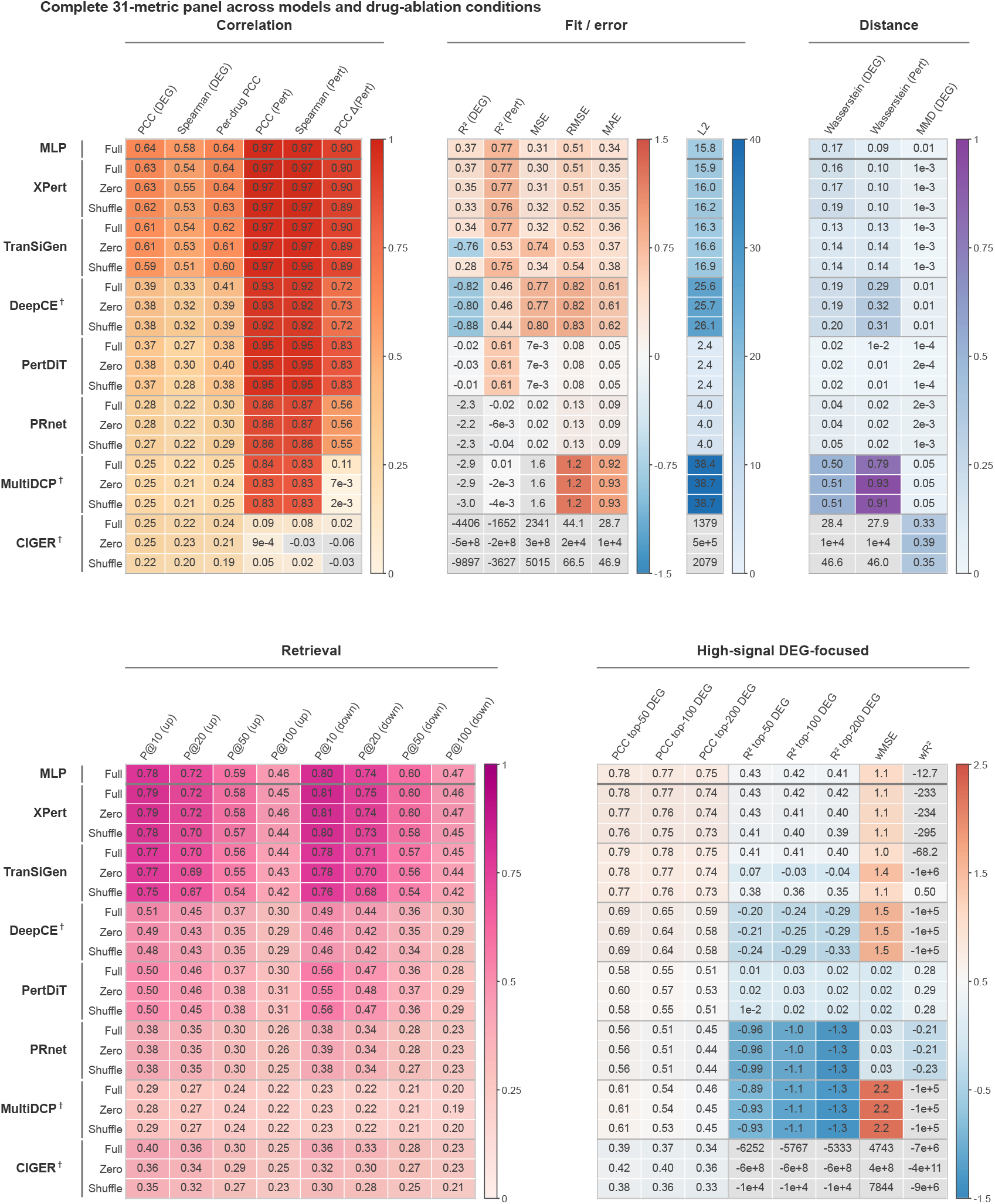
Complete 31-metric panel across models and drug-ablation conditions. Heatmap of 31 evaluation metrics computed for the MLP (no drug) and the seven published models under the Full, Zero and Shuffle input conditions. Metrics are organised into five evaluation families: Correlation, Fit/Error, Distance, Retrieval and High-signal DEG-focused metrics. Cell colour encodes the across-fold mean on a family-specific scale; colour scales are intended to show within-family patterns rather than direct cross-family magnitude comparisons. Darker colours indicate better performance within each metric family. † Models not originally designed to accept sample-paired L1000 basal expression profiles.

**Extended Data Fig. 5:**
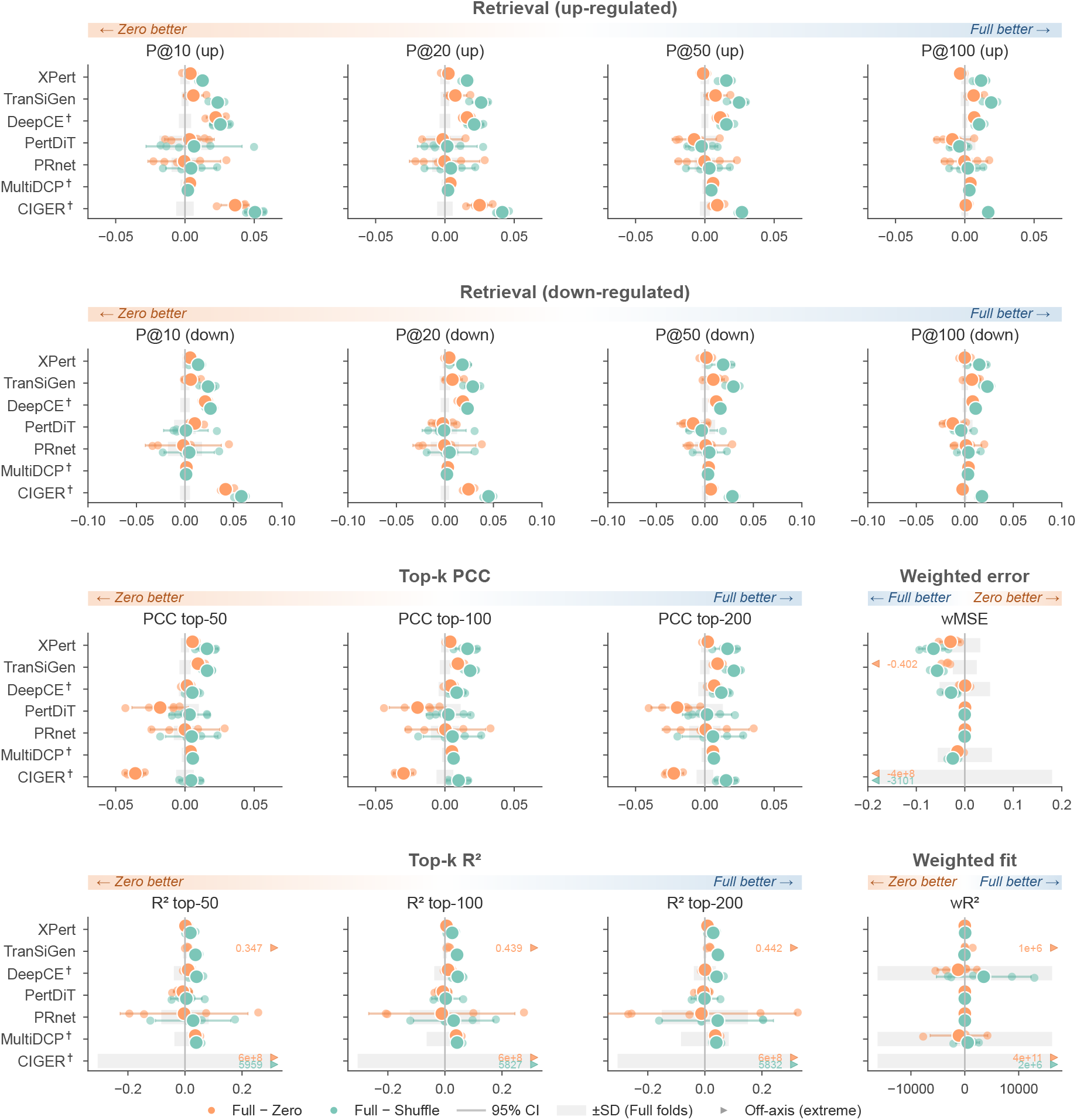
Drug-ablation effect sizes across signal-focused metrics. Paired drug-ablation effect sizes across signal-focused metrics. For each model and metric, orange points show Δ = Full −Zero and teal points show Δ = Full − Shuffle. Signal-focused metrics include retrieval of top-ranked up- and down-regulated genes, top-*k* PCC, top-*k R*^2^, weighted error and weighted fit metrics. Points denote across-fold means and horizontal segments denote 95% CIs across five folds (Student’s *t*, df = 4); vertical dashed lines indicate Δ=0. Grey bands indicate *±*s.d. across Full-condition folds. Direction bands indicate the favourable direction for each metric, and triangles or external labels denote off-axis extreme values. † Models not originally designed to accept sample-paired L1000 basal expression profiles.

**Extended Data Fig. 6:**
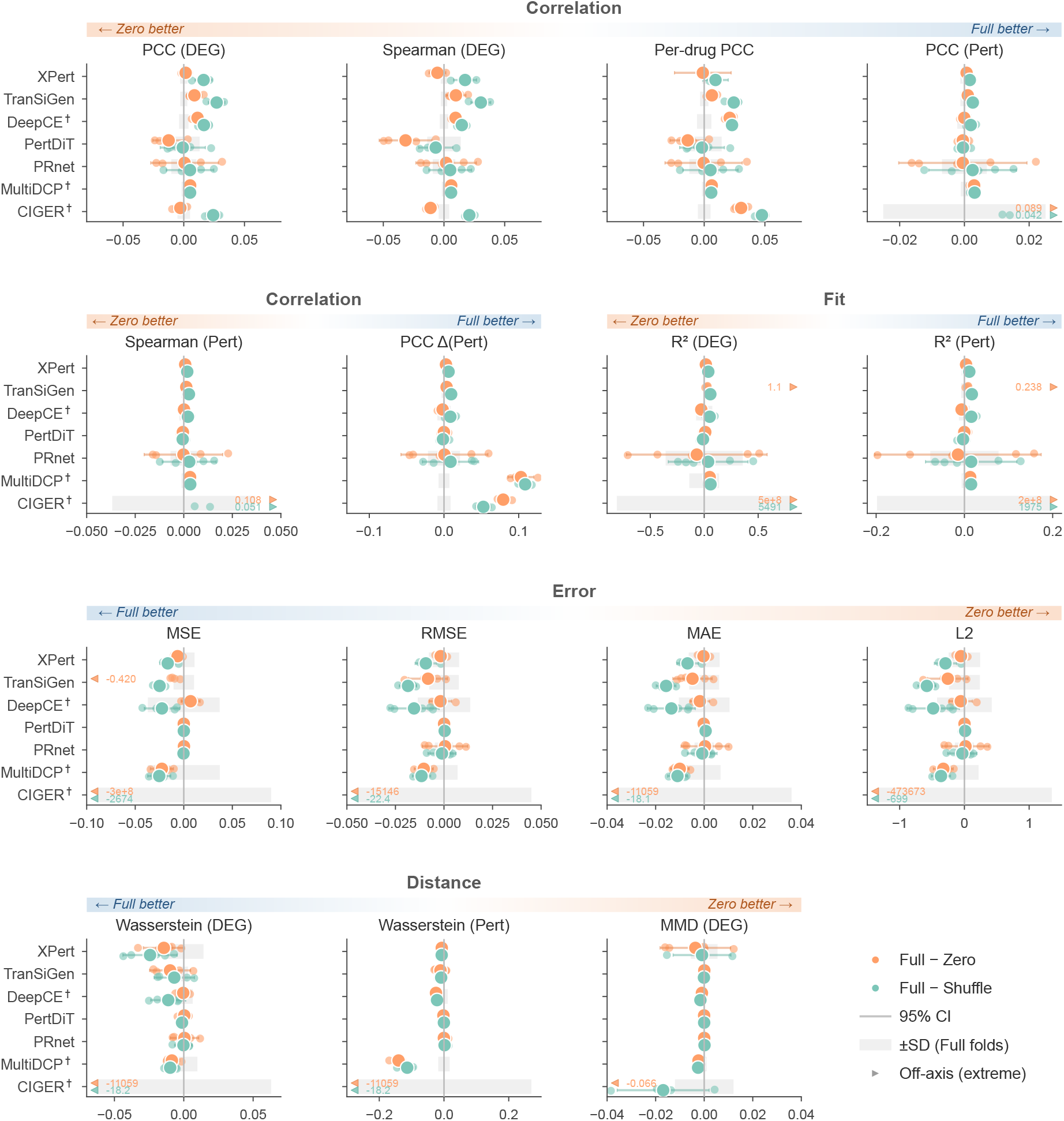
Drug-ablation effect sizes across core metrics. Paired drug-ablation effect sizes across core performance metrics. For each model and metric, orange points show Δ = Full −Zero and teal points show Δ = Full −Shuffle. Core metrics include correlation, fit, error and distributional-distance measures. Points denote across-fold means and horizontal segments denote 95% CIs across five folds (Student’s *t*, df = 4); vertical dashed lines indicate Δ=0. Grey bands indicate *±* s.d. across Full-condition folds. Direction bands indicate the favourable direction for each metric, and triangles or external labels denote off-axis extreme values. † Models not originally designed to accept sample-paired L1000 basal expression profiles.

**Extended Data Fig. 7:**
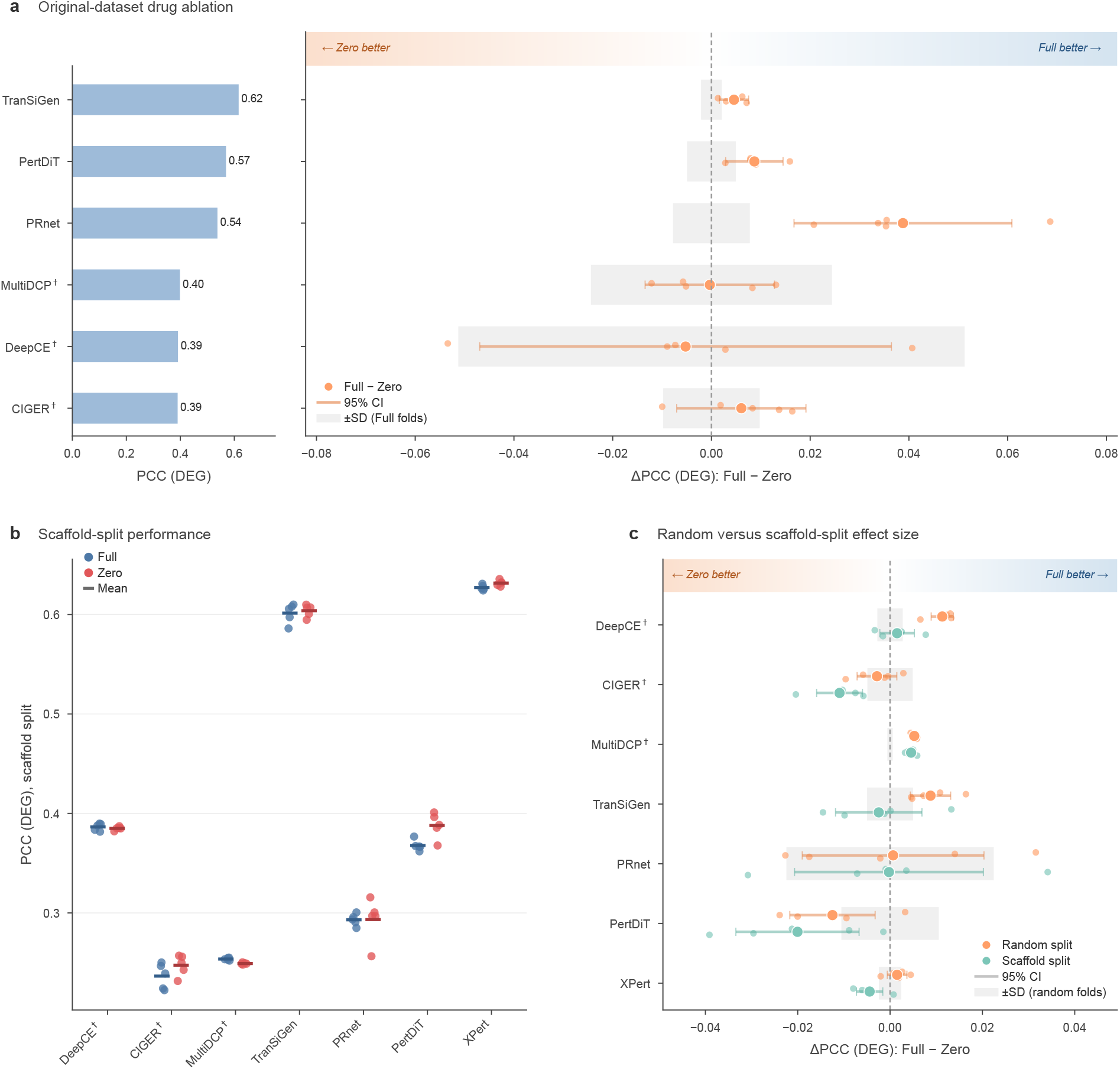
Drug-ablation effects across original and scaffold-based evaluation settings. **a**, Drug-ablation performance on each model’s original source task. Left, Full-condition performance, shown as PCC (DEG) where directly computable. Right, paired effect size ΔPCC (DEG) = Full − Zero. Models were evaluated on their original published datasets and splits where available, with drug-specific representations replaced by zeros at the architectural intervention points shown in Extended Data Fig. 1. **b**, Scaffold-split PCC (DEG) under Full and Zero conditions. Test compounds were assigned to folds by Bemis–Murcko scaffold. Points indicate fold-level estimates and horizontal lines indicate across-fold means. **c**, Paired effect size ΔPCC (DEG) = Full − Zero compared between random drug-blind and scaffold-based splits. Points denote across-fold means and horizontal segments denote 95% CIs across five folds (Student’s *t*, df = 4). Grey bands indicate *±* s.d. across random-split folds. Positive values indicate better performance under the Full condition. † Models not originally designed to accept sample-paired L1000 basal expression profiles.

**Extended Data Fig. 8:**
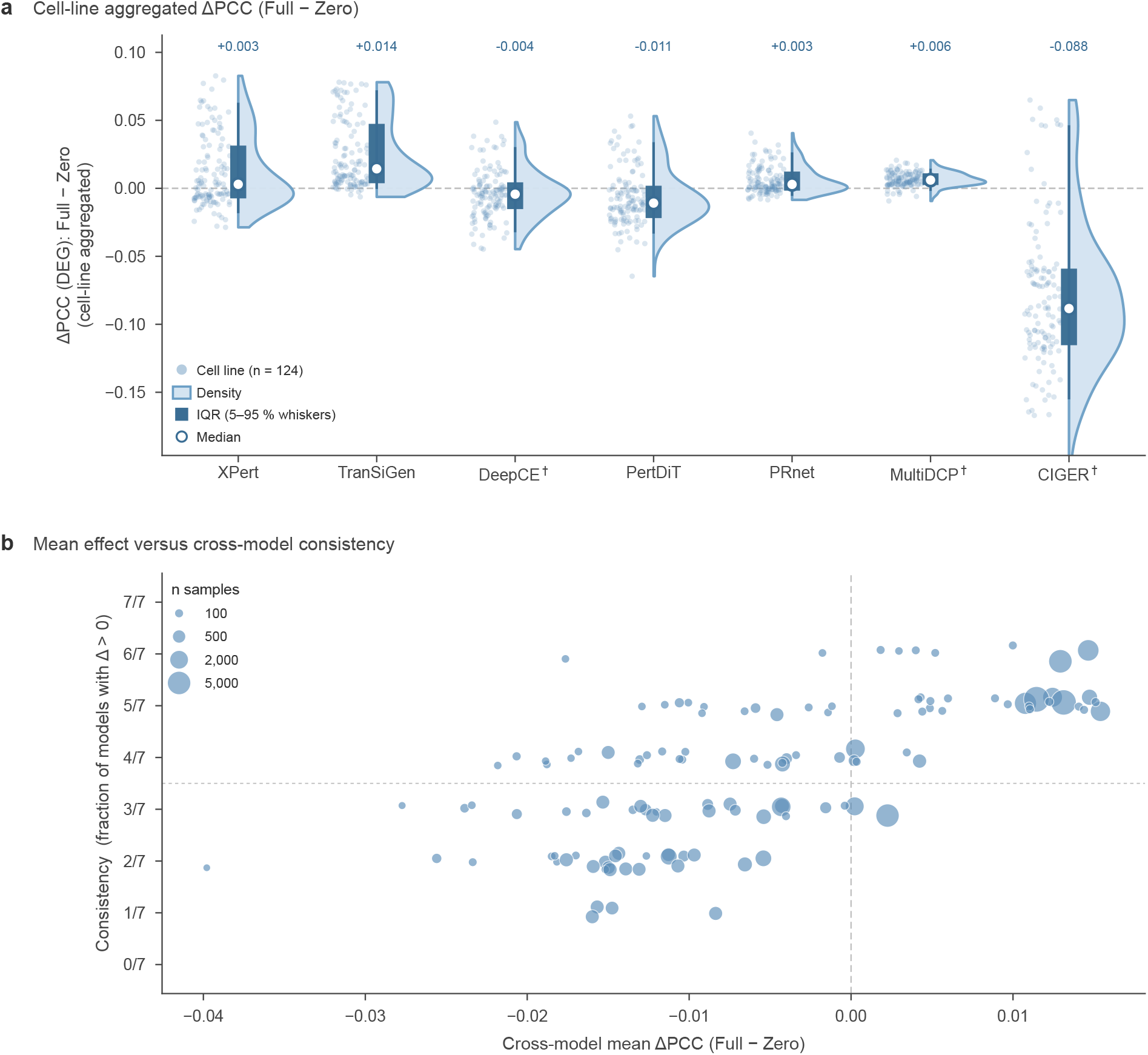
Cell-line level heterogeneity of drug-ablation effects. **a**, Cell-line aggregated ΔPCC (DEG), computed as PCC (DEG)_Full_ − PCC (DEG)_Zero_, for each model. Each point represents a cell line, half-violins show the empirical density, boxes show the interquartile range with 5–95% whiskers and open circles indicate medians. The dashed horizontal line marks Δ = 0. Values above each model indicate the median cell-line aggregated ΔPCC. **b**, Cross-model consistency of cell-line drug-ablation effects. Each point represents one cell line; the *x* axis shows the mean ΔPCC across models and the *y* axis shows the fraction of models for which Δ *>* 0. Bubble area is proportional to the cell line’s total test-sample count. † Models not originally designed to accept sample-paired L1000 basal expression profiles.

**Extended Data Fig. 9:**
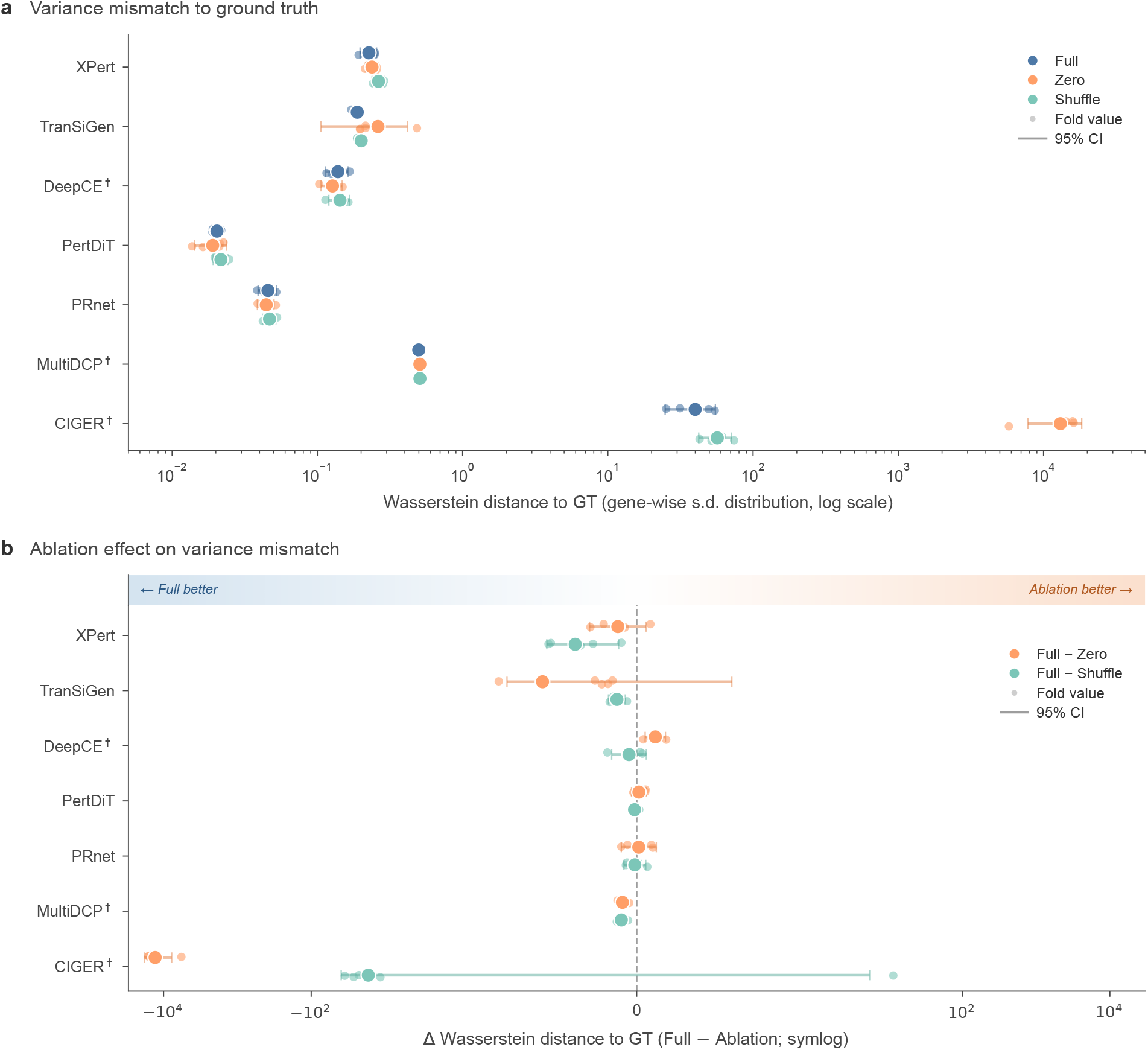
Variance calibration under drug ablation. **a**, Wasserstein distance between the predicted and ground-truth gene-wise s.d. distributions for each model under the Full, Zero and Shuffle conditions. Distances are shown on a log scale; lower values indicate closer agreement with the ground-truth variance distribution. Small points indicate fold-level estimates, larger points indicate across-fold means and horizontal segments indicate 95% CIs across five folds (Student’s *t*, df = 4). **b**, Ablation effect on variance mismatch, computed as ΔWasserstein = Wasserstein(Full) − Wasserstein(ablation) for the Zero and Shuffle ablations. The *x* axis is shown on a symlog scale and the vertical dashed line marks Δ = 0. Negative values indicate that the Full condition is closer to the ground truth, whereas positive values indicate that the ablated condition is closer to the ground truth. † Models not originally designed to accept sample-paired L1000 basal expression profiles.

**Extended Data Fig. 10:**
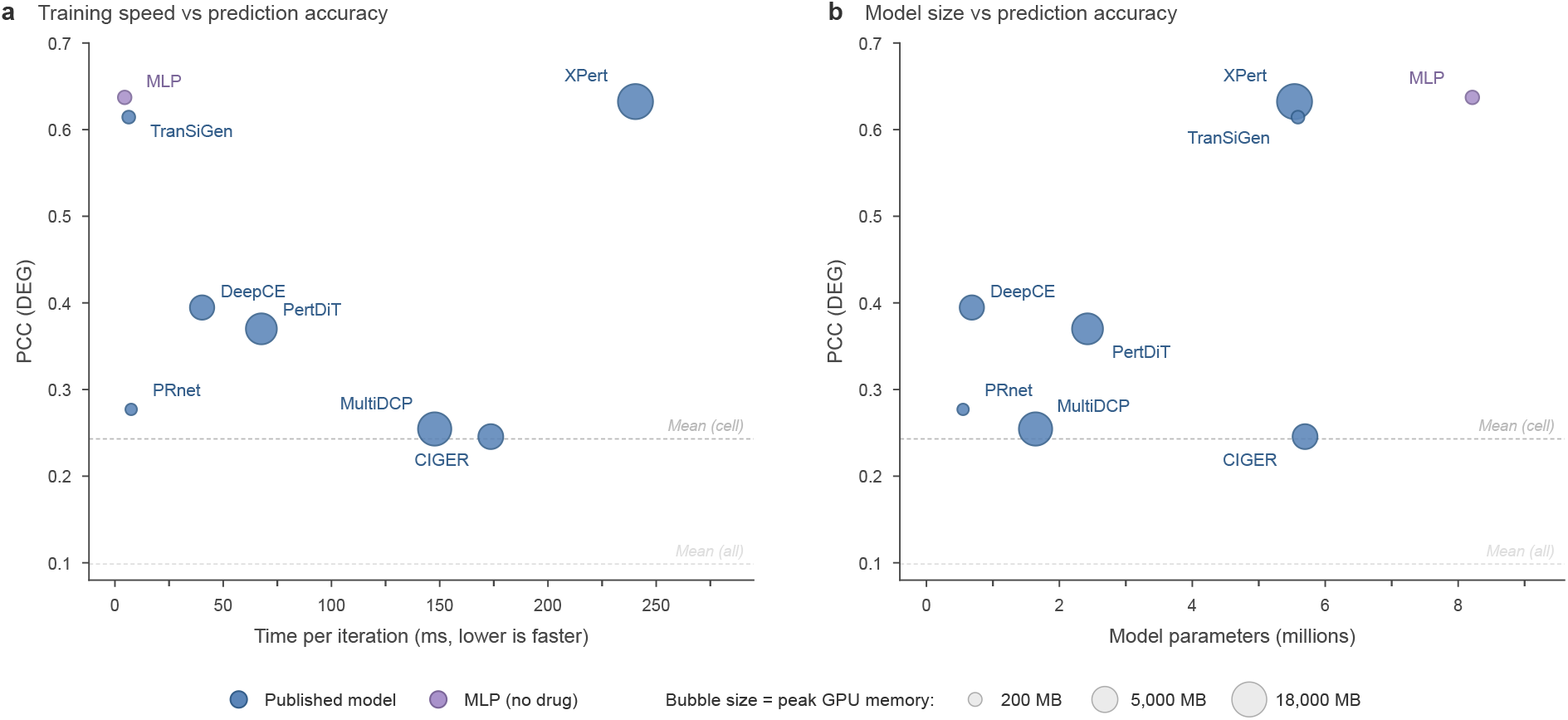
Computational cost versus prediction accuracy. **a**, Training speed, measured as milliseconds per iteration, versus mean drug-blind PCC (DEG) for the seven published models and the MLP (no drug). **b**, Number of trainable parameters versus mean drug-blind PCC (DEG) for the same methods. Bubble area is proportional to peak GPU memory during training. Blue points denote published models and the purple point denotes the MLP (no drug). Dashed horizontal lines indicate the Mean (cell) and Mean (all) baselines. Efficiency measurements were performed under the same measurement protocol.

